# The role of methylation and structural variants in shaping the recombination landscape of barley

**DOI:** 10.1101/2024.07.22.604552

**Authors:** Federico Casale, Christopher Arlt, Marius Kühl, Jinquan Li, Julia Engelhorn, Thomas Hartwig, Benjamin Stich

## Abstract

Meiotic recombination is not only a key mechanism for sexual adaptation in eukaryotes but crucial for the accumulation of beneficial alleles in breeding populations. The effective manipulation of recombination requires, however, a better understanding of the mechanisms regulating the rate and distribution of recombination events in genomes. Here, we identified the genomic features that best explain the recombination variation among a diverse set of segregating populations of barley at a resolution of 1 Mbp and investigated how methylation and structural variants determine recombination hotspots and coldspots at a high-resolution of 10 kb. Hotspots were found to be in proximity to genes and the genetic effects not assigned to methylation were found to be the most important factor explaining differences in recombination rates among populations along with the methylation and the parental sequence divergence. Interestingly, the inheritance of a highly-methylated genomic fragment from one parent only was enough to generate a coldspot, but both parents must be equally low methylated at a genomic segment to allow a hotspot. The parental sequence divergence was shown to have a sigmoidal correlation with recombination indicating an upper limit of mismatch among homologous chromosomes for CO formation. Structural variants (SVs) were shown to suppress COs, and their type and size were not found to influence that effect. Methylation and SVs act jointly determining the location of coldspots in barley and the weight of their relative effect depends on the genomic region. Our findings suggest that recombination in barley is highly predictable, occurring mostly in multiple short sections located in the proximity to genes and being modulated by local levels of methylation and SV load.

## INTRODUCTION

In sexual reproduction, the recombination between the maternal and paternal homologous chromosomes followed by their random assortment during meiosis generates new combinations of alleles that can be transmitted to the next generation (Barton and Charlesworth 1998). This mechanism for generating genetic diversity is widely conserved among eukaryotes since it provides a possibility for the adaptation of species: the reshuffling of alleles breaks the linkage between beneficial and deleterious mutations, allowing the accumulation of beneficial mutations into one haplotype, and, ultimately, generating new phenotypes upon which selection can act (Muller 1932; Peck 1994).

Meiosis consists of a single round of DNA replication in which the bivalent is generated –a pair of physically linked homologous chromosomes each composed of two replicated sister chromatids– followed by two rounds of chromosome segregation (for a review see Mercier et al. 2015). During the first round, meiotic recombination occurs in prophase I when programmed DNA double-strand breaks (DSBs) are repaired as either crossovers (COs), the reciprocal exchange of large regions between homologous nonsister chromatids, or noncrossovers (NCOs), –the unidirectional copies of small fragments from any of the intact homologous chromatids (Szostak et al. 1983). In addition, when both CO and NCO occur, mismatches at the site of the strand invasion are produced if sequence polymorphisms exist among homologous chromosomes. Such mismatches are either restored to the original allelic state (i.e., intersister repair) or repaired in favor of the homologous allele resulting in gene conversion (GC): the nonreciprocal exchange of alleles between homologous nonsister chromatids (Burt 2000; Wijnker et al. 2013).

The rate of recombination events is strongly regulated and has been proposed to be related to an optimal level of recombination for adaptation in eukaryotes (Mercier et al. 2015). The presence of at least one CO per bivalent, termed the obligate CO, is required for the correct segregation of homologous chromosomes (Hall 1972). In addition, the number of generated DSBs exceeds the number of observed COs which are rarely more than three per chromosome per meiosis in most species (Martini et al. 2006; Baudat and Massy 2007; Bauer et al. 2013). Most of the observed COs, furthermore, prevail in small regions of a few kilobases called recombination hotspots where CO rates are several times greater than the chromosome average (Mézard 2006; Choi and Henderson 2015). Remarkably, both the rate and distribution of COs in the genome have been shown to exhibit extensive inter- and intraspecific variation (Nachman 2002; Ritz et al. 2017; Lawrence et al. 2017). The mechanisms behind such variation, however, are not completely understood. Such knowledge is required for the effective manipulation of recombination, e.g., for the purpose of plant breeding. This recombination determines the frequency of breaking undesirable linkages and stacking favorable alleles in the genetic background of breeding populations and defines marker resolution to map quantitative traits (Bauer et al. 2013; Blary and Jenczewski 2019).

Earlier studies provided valuable information for deciphering the mechanisms behind recombination variation in plants. Most of the recombination in plants occurs in euchromatic regions where chromatin is accessible while heterochromatin is suppressed for meiotic recombination (Henderson 2012). For example, earlier studies in *Arabidopsis thaliana* and other species have shown that recombination events tend to occur in genomic regions with hypomethylated DNA (Yelina et al. 2012; Rodgers-Melnick et al. 2015; Marand et al. 2019; Apuli et al. 2020; Fernandes et al. 2024) and depleted nucleosome density (Choi et al. 2013; Wijnker et al. 2013). Moreover, COs in plants are typically located in close proximity to genes and in association with chromatin marks that favor transcription (Choi et al. 2013; Mercier et al. 2015). A positive correlation between the recombination rate and gene density has been observed in many plant families (Paape et al. 2012; Choi et al. 2013; Silva-Junior and Grattapaglia 2015; Gion et al. 2016; Wang et al. 2016; Apuli et al. 2020), including most grasses (Bauer et al. 2013; Darrier et al. 2017; Jordan et al. 2018; Gardiner et al. 2019; Marand et al. 2019; Casale et al. 2022). It is worth noting that some studies have instead reported a negative correlation between recombination rate and gene density (Kim et al. 2007; Giraut et al. 2011; Yang et al. 2012; Rodgers-Melnick et al. 2015).

Polymorphisms among homologous chromosomes are expected to prevent recombination due to defective strand invasion and homology pairing caused by the increase in mismatches among nonsister chromatids (Henderson 2012). Earlier studies in plants, however, reported constrasting correlations between recombination rate and parental sequence divergence (Saintenac et al. 2011; Salomé et al. 2012; Yang et al. 2012; Bauer et al. 2013; Jordan et al. 2018; Marand et al. 2019). Recent reports in *Arabidopsis* suggested that the recombination rate has a positive correlation with parental allelic divergence until a level of mismatch among homologous chromosomes prevents CO formation (Blackwell et al. 2020; Hsu et al. 2022). Accordingly, large structural variants (SVs) were shown to suppress local recombination in several plant species (Rodgers-Melnick et al. 2015; Shen et al. 2019; Rowan et al. 2019; Fernandes et al. 2024).

The above-mentioned contrasting findings impose the necessity to characterize the associations between recombination and genomic features at the species level to avoid making incorrect assumptions. In addition, due to differences in the employed research methods, disagreements were observed among studies on the same species (Apuli et al. 2020). In this respect, most reported recombination rates in plants were calculated based on a coarse genomic resolution that failed to capture the complete genetic variation generated from meiosis. At present, most reported recombination assessments at high-resolution have been performed in *Arabidopsis* (Sun et al. 2012; Lu et al. 2012; Yang et al. 2012; Wijnker et al. 2013; Rowan et al. 2019; Fernandes et al. 2024), while only a few have been performed in major cereal crops such as maize (*Zea mays*) (Rodgers-Melnick et al. 2015; Li et al. 2015), rice (*Oryza sativa*) (Si et al. 2015; Marand et al. 2019), and wheat (*Triticum aestivum*) (Jordan et al. 2018). In barley (*Hordeum vulgare*), the fourth of this list, earlier low-resolution recombination studies successfully revealed the genetic basis of recombination as well as the association of recombination with some environmental and genomic features on a broad genomic scale (Higgins et al. 2012; Dreissig et al. 2019, 2020; Casale et al. 2022); however, a high-resolution study depicting the complete meiotic recombination landscape and the respective associations of genetic and epigenetic features in the barley genome is still lacking.

Consequently, in the present study, we aimed to (*i*) identify the genomic features that best explain the recombination variation among the double round-robin (DRR) populations, (*ii*) detect recombination events in the barley genome at high-resolution, and (*iii*) analyze the effect of genomic features in determining the location of recombination hotspots and coldspots in the genome.

## RESULTS

### The genomic features associated with recombination rate variation in the barley genome

The recombination rate was almost null in the pericentromeric region, but increased toward the distal regions of the chromosomes (Figure 1A). The SV load, the physical fraction spanned by the analyzed SVs (insertions, deletions, inversions, duplications, and translocations) increased toward the distal regions of the chromosomes as did the recombination rate and gene density (Figures 1A and S1). In this way, the presence of SVs and the sequence divergence among parents showed a significant positive Pearson’s correlation (P *<* 0.05) of 0.35 with the recombination rate along the distal regions of the chromosomes (Figures 1C and S3A), whereas a lower but significant correlation (P *<* 0.05) of 0.1 was observed in the pericentromeric region (Figures 1D and S3B). The sequence divergence among the parental inbred lines, which showed a significant positive correlation (P *<* 0.05) with the SV load of 0.27, was found to have a significant positive correlation (P *<* 0.05) with recombination rates of only 0.1 and 0.14 in the distal and pericentromeric regions, respectively (Figures 1A, 1C, and 1D). The methylation level in the sequence contexts differed along the barley chromosomes, where the level in the CpG and CHG contexts reached a maximum in the pericentromeric region and decreased toward the telomeres, while the methylation level in the CHH context increased toward the telomeres (Figure 1B). Because the CpG and CHG contexts had higher overall methylation levels than did the CHH context, the average methylation level along the chromosomes mostly represented the trend observed for the CpG and CHG contexts. The observed significant negative correlation (P*<* 0.05) between the average methylation level and recombination rate along the barley chromosomes was therefore due to the CpG and CHG contexts but not to the CHH context (Figures 1C and 1D). The difference in the methylation level between the parental inbred lines of any of the analyzed populations was greater in the distal regions than in the pericentromeric regions of the chromosomes in all three analyzed sequence contexts (Figure S4). This explained the positive correlation (P*<* 0.05) of 0.17 of such difference and the recombination rate along the distal regions in the barley chromosomes (Figures 1C, 1D, S3A, and S3B).

**Fig. 1:**
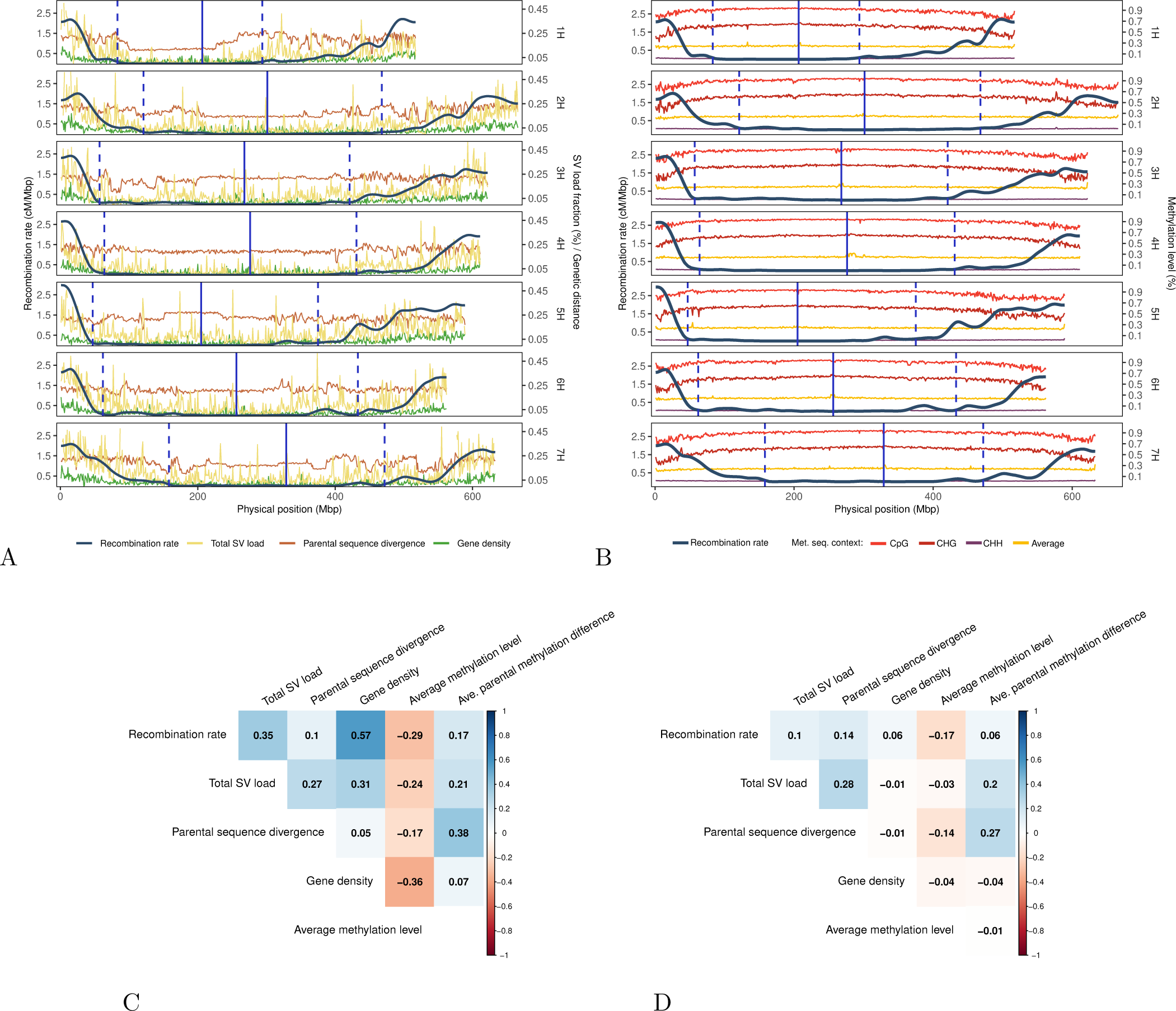
(A) Distribution of recombination rate, total structural variants’ load fraction, gene density, and parental sequence divergence between the respective parental inbred lines on average across the 45 double round-robin (DRR) populations in 1 Mbp windows across the seven barley chromosomes. The total structural variants (SV) load fraction represents the portion spanned of a given 1 Mbp genomic window by the sum of insertions, deletions, invertions, and duplications. (B) Distribution of recombination rate and methylation level in 1 Mbp windows for the sequence contexts CpG, CHG, CHH, and their mean, on average across the seven barley chromosomes and 45 DRR populations. The methylation level values for a given methylation sequence context represent the percentage of methylated reads of such context present in a 1 Mbp window averaged across the 45 DRR populations. The vertical blue solid line indicates the position of the centromere in the Morex v3 reference genome and the vertical blue dashed lines indicate the pericentromeric region calculated across the 45 DRR populations. (C-D) Correlation matrix between recombination rate, total SV load fraction, parental sequence divergence, gene density, average methylation level, and parental difference in methylation level, across 1 Mbp windows in the distal (C) and the pericentromeric (D) regions of the barley chromosomes for the average across the 45 DRR populations. Pearson’s correlation coefficients are indicated with a color gradient from −1 (red) to 1 (blue).

### The genomic features associated with recombination rate variation among barley populations

The best subset of genomic features explaining the recombination rate variation among the 45 DRR populations in 1 Mbp genomic windows along the barley chromosomes was identified using a stepwise regression model (Figure 2). The fraction of 1 Mbp windows of the barley genome in which a given genomic feature was found to be significantly correlated and the direction of such correlation are given in parentheses below. The genetic effects, which were calculated from the GRE, were found to be the most determining factor in explaining differences in recombination rate among populations with a promoting effect on both the distal (positive, 0.76) and the pericentromeric regions (positive, 0.40) of the chromosomes. The parental sequence divergence was also found to be positively correlated with the recombination rate in both mentioned chromosomal regions (positive, 0.24 and 0.19, respectively). A few windows showed a significant negative correlation between parental sequence divergence and the recombination rate (negative, 0.04 and 0.01 in both respective chromosomal regions). Notably, the windows with a negative correlation had a parental sequence divergence near its relative maximum, which appeared to be associated with a high SV load (Figure S5A). The average methylation level across the different sequence contexts was negatively correlated with the recombination rate in the distal (negative, 0.43) and pericentromeric (negative, 0.12) regions of the barley chromosomes. This means that populations with a higher methylation level than others in a particular window showed a lower recombination rate than others in that window, and vice-versa. Additonally, the multiple regression model used to calculate genetic effects revealed a correlation between the methylation level and the sum of the GREs ranging from *−*0.29 to 0.03 per 1 Mbp genomic window with an average of *−*0.06 in the distal region. The difference in the average methylation level between the parental inbred lines of the respective populations was found to have only a low impact on the recombination rate variation among populations (in distal regions, positive, 0.04; negative, 0.03). The physical fraction of 1 Mbp genomic windows spanned by all SVs was found to have a low (and mostly positive) impact on the differences in recombination rate among populations (in distal regions, positive, 0.05; negative, 0.02).

**Fig. 2:**
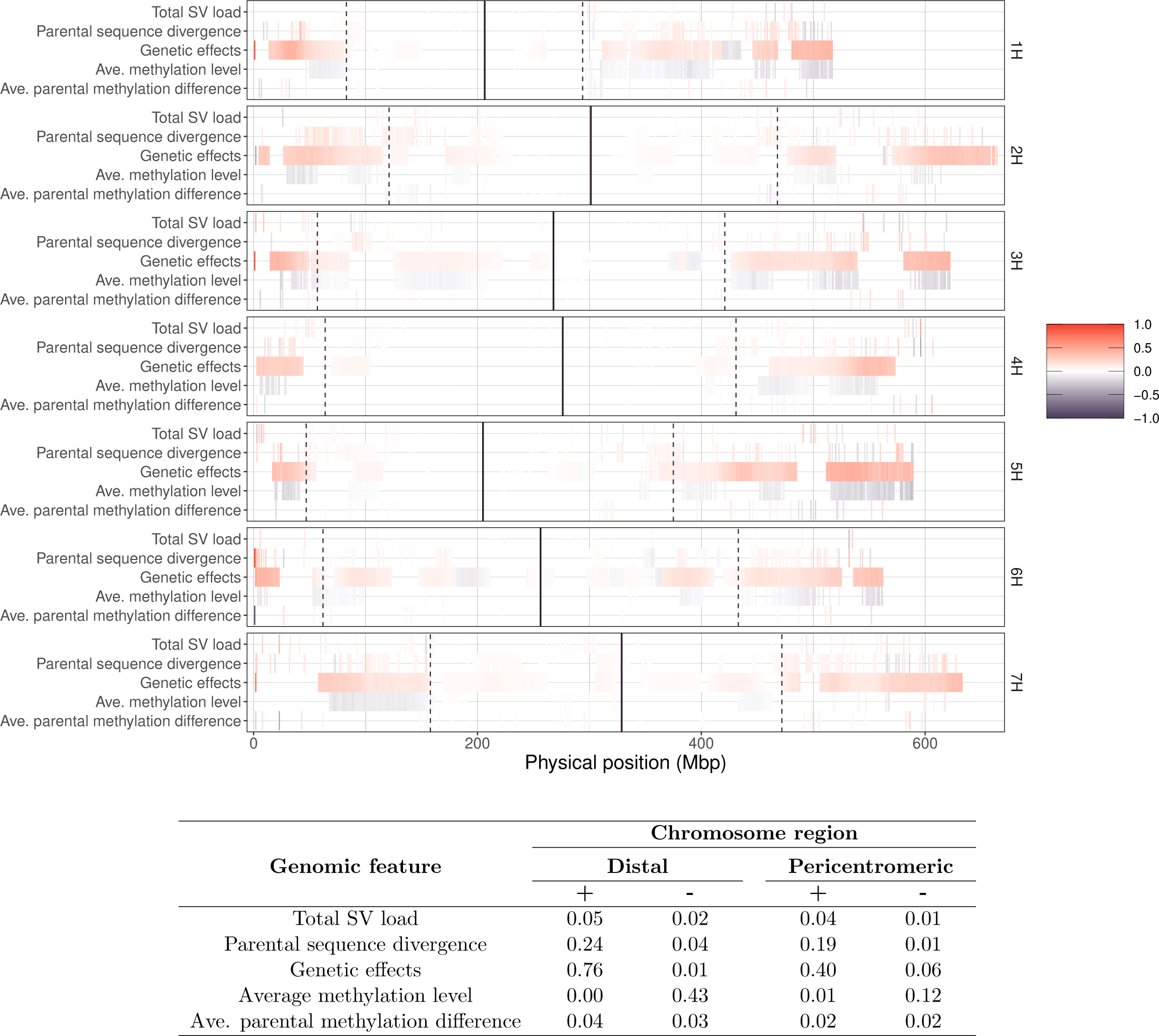
The genomic features explaining the variation in the recombination rate among the 45 double round-robin (DRR) populations in 1 Mbp windows across the seven barley chromosomes. The genomic features were selected for each window by a stepwise regression procedure. The standarized regression coefficients are indicated by a color gradient from −1 (purple) to 1 (red). The studied genomic features included sequence divergence among parental inbred lines, genetic effects, total structural variants (SV) load, average methylation level across the sequence contexts CpG, CHG, and CHH, and the difference in the methylation level among the parental inbred lines for the average of such sequence contexts. The vertical solid line indicates the position of the centromere in the reference genome, and the vertical dashed lines indicate the pericentromeric region calculated across the 45 DRR populations. The analysis was performed separately for the distal and pericentromeric regions of the barley chromosomes, respectively. The fraction of 1 Mbp windows of the barley genome in which the genomic features were found to be significantly associated with the recombination rate variation across the 45 DRR populations are displayed in the table. The fractions of 1 Mbp windows positively and negatively associated with the recombination rate were indicated with the “+” or the “*−*” signs, resdectvely.

The methylation level in the three sequence contexts –CpG, CHG, and CHH– analyzed independently in an extended model was shown to have the same repressive effect on recombination as the average across the three sequence contexts (Figure S6 and Table S2). In addition, the parental methylation difference for the sequence contexts CHG and CHH was found to be positively associated with the recombination rate, and the inverse was found for the context CpG. Furthermore, the extended model revealed no impact of the different SV types on recombination rate variation among populations.

### The key parameters for detecting recombination events at high-resolution

The mRNA sequencing of recombinant inbred lines (RILs) of the evaluated populations yielded 4.1 M sequence variants of which, 858 K SNPs remained after being intersected with reported SNV parental data generated by DNA sequencing. A total of 12 K SNPs were added from the iselect array, resulting in a total of approximately 870 K SNPs genome-wide. After reoving SNPs carrying nonparental alleles or with 100% missing data, the final number of genome-wide SNPs ranged from 214 to 259 K per population (Table S3 and Figure S7). The median inter-SNP distance was 132 bp on average across populations, which was 79 times shorter than the 10, 475 bp utilized to count COs for the same three populations analyzed in a previous study (Casale et al. 2022). The maximum inter-SNP distance ranged from 5.45 to 11.99 Mbp among populations (Table S3), denoting large regions that were identical by descent (IBD) among the parental inbred lines involved in a given population and thus among their respective offspring. This resulted in an average density of 57 SNPs per Mbp across populations. The mean length for each parental SNP block category was 14.8, 44.1, and 111.5 Mbp for the short, medium, and long CO-related blocks, respectively (Table S4). The block length was positively correlated with the number of SNPs per block (*>* 0.75, *P <* 0.001). Therefore, the false positive rate for detecting CO was inversely proportional to the marker block length, supporting the identification of block length categories with different CO layers.

On average, there were were 30, 87, and 269 genome-wide COs accumulated per RIL across populations for the long, medium, and short block lengths, respectively (Table S5). Considering only the layer of COs generated by blocks longer than 3 Mbp, the genome-wide CO counts per RIL ranged from 14 to 65 across populations (Figures S8 and S9). Since a given CO breakpoint was determined as the midpoint of the CO interval, the breakpoint location accuracy depended on the CO interval length (Table S6). The lengths of the detected CO intervals ranged from less than 20 bp to 10.8 Mbp with a median that varied from 41.9 kb to 151.6 kb depending on the considered CO layer. The average number of genome-wide GC events that accumulated per RIL across populations was 58, 251, and 6,521 for long, medium, and short GC-related block lengths, respectively.

An average of 80 recombination hotspot windows per chromosome were found across the three selected populations (Table S7). Among these, 12 were found in the pericentromeric regions, and the rest were found in the distal regions of the barley chromosomes (Table S8 and Figure S10). The recombination hotspot windows contained 0.26, 0.14, and 0.25 of the total observed COs in the HvDRR13, HvDRR27, and HvDRR28 populations, respectively. Less than 10% of the hotspot windows in a given population were shared with another population and less than 1% of the total counted hotspot windows were shared among the three analyzed populations (Figure S11). Interestingly, both the number and conservation level of coldspot windows far exceeded those of hotspots. On average, across populations, more than half (10,436 out of the 19,496) of the distal windows were recombination coldspots. More than 60% of the coldspot windows in a given population were shared with the other populations, and 16.7% of the coldspot windows were present in all three analyzed populations. The majority of the coldspot windows were located contiguously in regions with lengths that varied from 10 kb to 17 Mbp with an average of 322 kb across the three analyzed populations.

More than 15% of the GC hotspot windows in a given population were shared among the three analyzed populations (Figure S12). The GC hotspot windows were found to overlap with the CO hotspot windows in the distal region of the barley chromosome significantly more than they did under a random distribution across the three analyzed populations (Table S9). The GC hotspot windows detected in a given population overlapped with 12–15% of the CO hotspot windows in the same population and with 7.5–9.5% of the CO hotspot windows detected in the other two analyzed populations (Table S10). There was no significant (*P >* 0.05) difference between such overlap proportions in the HvDRR27 and HvDRR28 populations.

### The effect of methylation and structural variants on recombination rate variation at high-resolution The coldspot and hotspot windows have different methylation level and SV load than the rest of the genome

The coldspot and hotspot windows in a given chromosome region showed distinct methylation patterns compared to the average remaining windows in the same region in the three analyzed populations (Figure 3). The average methylation level across all three methylated sequence contexts in the coldspot windows of the distal telomeric subregion was significantly greater (*P <* 0.001) than that across the other windows in both distal subregions. In contrast, the coldspot windows in the distal proximal region were not found to be differentially methylated (*P >* 0.001) from other windows in any of the distal subregions. However, when analyzing the CpG and CHG sequence contexts separately, the methylation level in the coldspot windows of the distal proximal subregion was found to be significantly greater (*P <* 0.001) than that in the other windows of both distal subregions. Differently, the methylation level in the coldspot windows of the distal telomeric subregion was significantly greter (*P <* 0.001) than that in the other windows in this subregion but significantly lower (*P <* 0.001) than that in the windows of the distal proximal subregion. The coldspot windows in such comparisons that were below the critical value (methylation levels of 0.89 and 0.59 for the sequence contexts CpG and CHG, respectively) were found to have a significantly (*P <* 0.001) greater total SV load fraction than the coldspot windows above the critical value (Table S11).

**Fig. 3:**
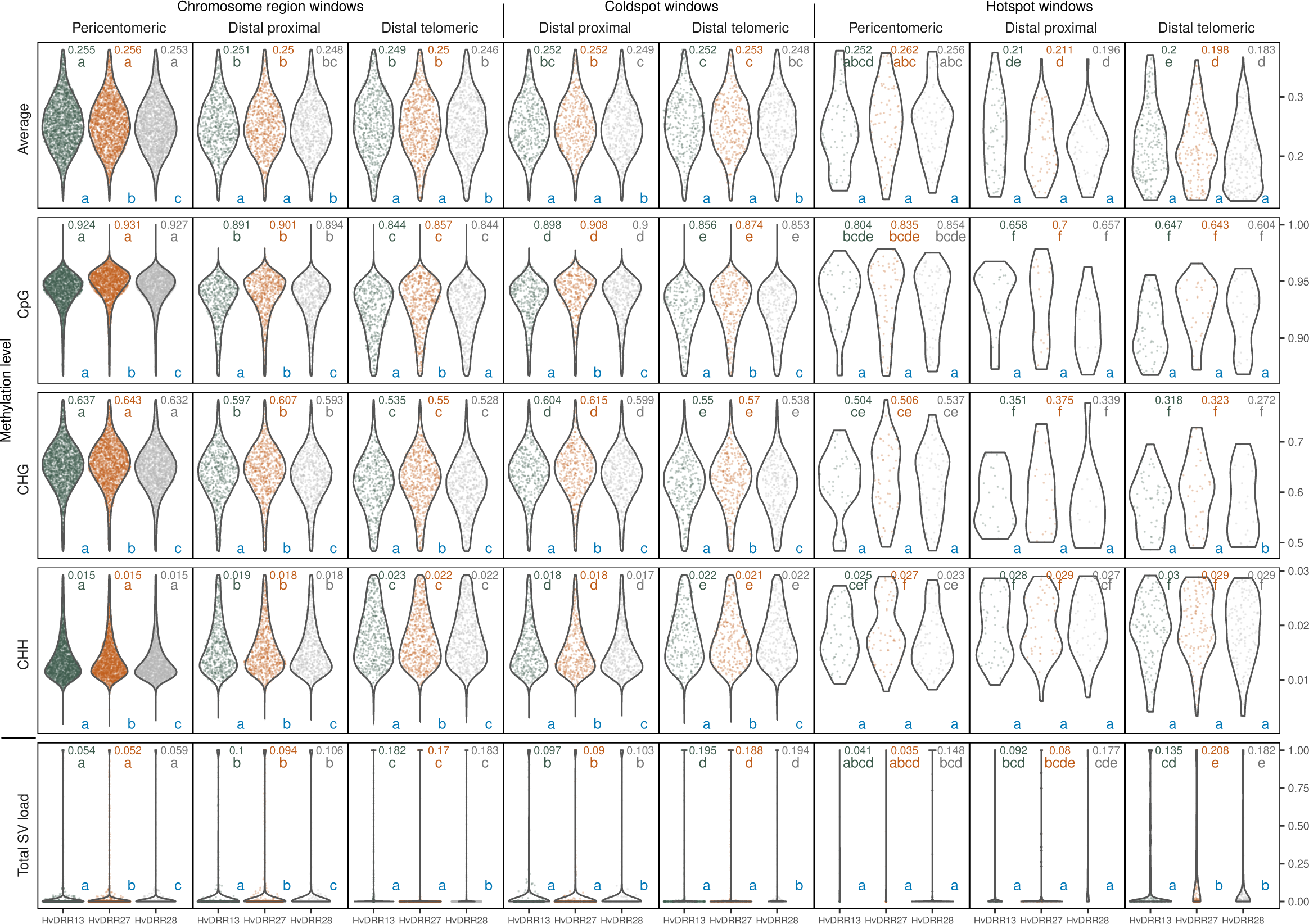
Distribution of the methylation level for the methylated sequence contexts CpG, CHG, CHH, and their average, and the total structural variants (SV) load fraction in 10 kb genomic windows grouped by their location in different chromosomal regions-pericentromeric, distal proximal, and distal telomeric-, and recombination rate-total, coldspots, and hotspots- of the barley chromosomes in the analyzed double round-robin (DRR) populations HvDRR13 (green), HvDRR27 (orange), and HvDRR28 (grey). For each DRR population, the methylation level at each sequence context in a given genomic window was calculated as the average among the respective parental inbred lines’ methylation level values for the methylated cytosine positions in that window, weighted by the number of methylated cytosine positions corresponding to each parent. The displayed dots for chromosome regions and coldspots are a random subset of 1% of the total windows in each window group. For each genomic feature, the distribution’s mean of each window group from a given population is indicated with the related population color at the top-right of the respective plot. Significant differences (*α* = 0.001) in the Wilcoxon’s rank sum test among such means across window groups is indicated with different letters below the respective mean and sharing the population color. For a given genomic feature in each window grouping category, significant differences (*α* = 0.008) in the Wilcoxon’s rank sum test among populations are indicated with different blue letters at the bottom of each panel.

The average methylation level across the three sequence contexts in the hotspot windows was significantly lower (*P <* 0.001) than that across the other windows in any region of the barley chromosomes. The hotspot windows in the pericentromeric region, however, were not found to be differentially methylated from the rest of the windows in such region or from the windows in other regions of the genome, including coldspots. However, by analyzing the methylated sequence contexts separately, the hotspot windows were found to be significantly less methylated (*P <* 0.001) in the CpG and CHG sequence contexts than in the total windows in the pericentromeric region.

The coldspot windows in the distal telomeric regions were found to have a significantly greater total SV load (*P <* 0.001) than the rest of the windows in that subregion. In contrast, the coldspot windows in the distal proximal region did not show such an increase in total SV load. However, the coldspot windows below the critical value (SV loads equal to 0.187, 0.174, and 0.187 for the HvDRR13, HvDRR27, and HvDRR28 populations, respectively) in such comparisons had a significantly increased methylation level (*P <* 0.05) compared to the windows above the critical value for the CpG and CHG sequence contexts (Table S12). The hotspot windows were not observed to have a significantly different (*P >* 0.001) span of total SVs compared to the total windows in their respective chromosome regions. However, the observed overlaps between CO intervals and insertions/deletions and duplications were found to be significantly less frequent (*P <* 0.001) than such overlaps under a random distribution of the COs and the respective SVs in the distal regions of the barley chromosomes in the three analyzed populations (Table S13). In the case of inversions, such a pattern was observed only for the HvDRR27 population. Moreover, the distance between the CO breakpoints and the closest SV of any type was significantly greater (*P <* 0.001) then the CO-SV distances expected by chance (Table S14).

### The genomic environment neighboring hotspot and coldspot windows

The 10 kb genomic windows adjacent to coldspot regions were found to have a significantly lower (*P <* 0.001) average methylation level across the three sequence contexts than coldspots in both distal subregions of the chromosome in the three analyzed populations (Figure 4). This observation reflected the pattern produced at the methylated sequence contexts CpG and CHG, individually (Figure S13). In addition, any 10 kb window in the considered range from −40 kb to +40 kb around coldspot regions was found to have a significantly lower (*P <* 0.001) total SV load than the neighboring coldspot. The 10 kb genomic windows adjacent to hotspot regions were not found to have significantly (*P >* 0.001) different methylation levels or SV loads than any of the analyzed chromosomal regions or populations.

**Fig. 4:**
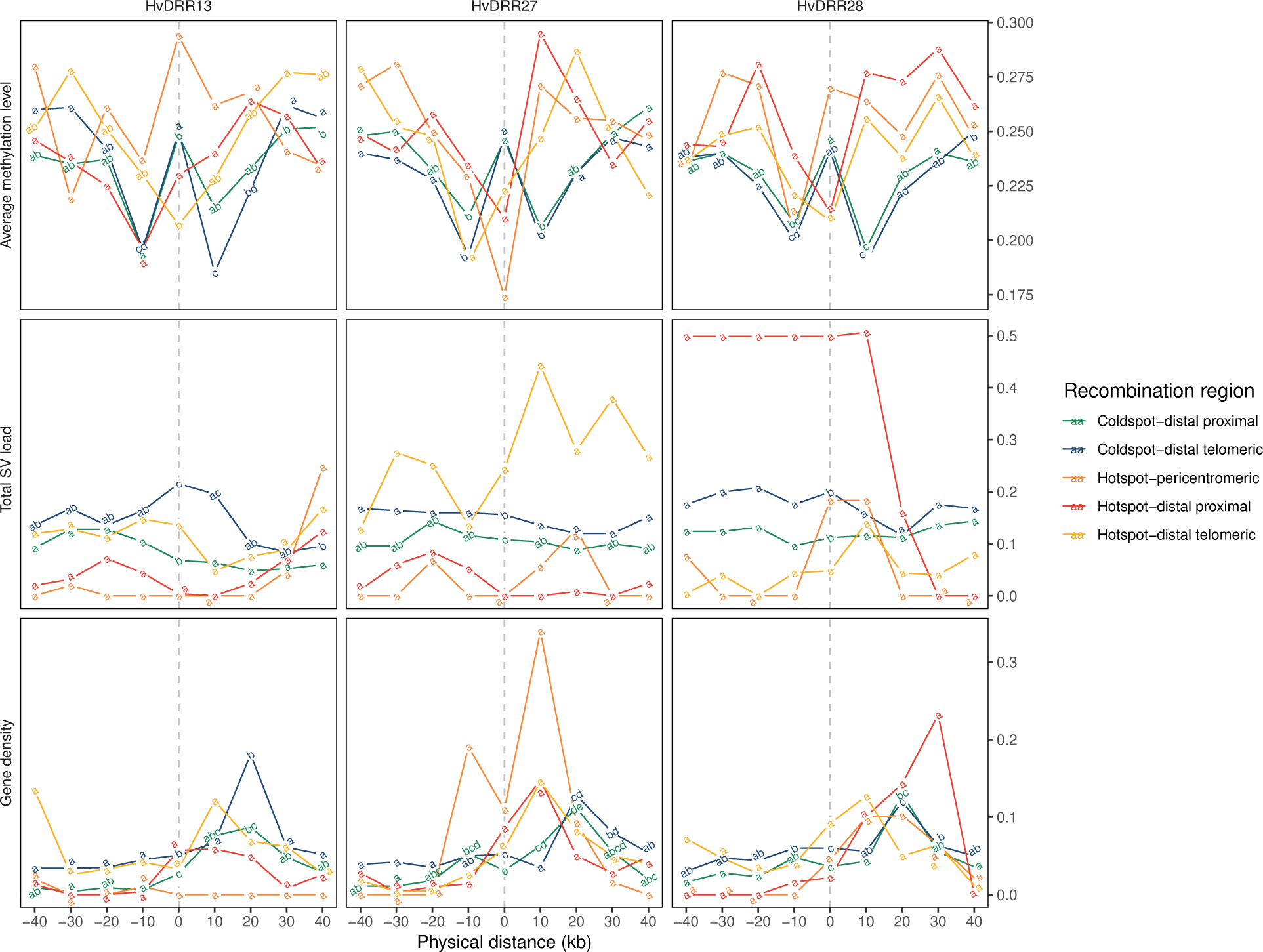
The mean values of the average methylation level among the methylated sequence contexts CpG, CHG, and CHH, total structural variants (SV) load, and gene density in the genomic windows grouped by the physical positions in the range from −40 kb to +40 kb around the coldspot and hotspot genomic regions in the pericentromeric, distal proximal, and distal telomeric region of the chromosomes in the analyzed double round-robin (DRR) populations HvDRR13, HvDRR27, and HvDRR28. The vertical dashed lines indicate the relative location of either the coldspot or the hotspot windows, respectively. The significant difference (*α* = 0.001) in the Wilcoxon’s rank sum test among the window groups corresponding to the different 10 kb physical position neighboring the respective hotspot or coldspot region in a particular chromosome region is indicated with different letters.

The windows neighboring coldspot regions were found to have a significantly lower (*P <* 0.001) gene density than these regions, except for the windows located 20 kb upstream of coldspots. However, the overlap between the coldspot regions and genes in the distal regions of the barley chromosomes was not significantly different (*P >* 0.001) from such overlap under a random distribution (Table 1). In contrast, a visual increase in the gene density from the hotspots to 20 kb upstream was observed in all the analyzed genomic regions, although this pattern was not significant (*P >* 0.001). Furthermore, the overlap between the hotspot regions and genes was found to be significantly (*P >* 0.001) greater than expected under a random distribution in the three analyzed populations, while the overlap between hotspots and intergenic regions was not found to be significantly (*P <* 0.001) greater than random, with the exception of the HvDRR27 population. In addition, a high proportion of the windows surrounding hotspot regions in both the proximal (0.37–0.49) and telomeric (0.33–0.45) subregions of the distal region of the genome were coldspot windows (Table S15).

**Table 1:** The observed overlaps among the recombination coldspot and hotspot 10 kb windows with genes and the intergenic regions in the distal region of the barley chromosomes, and their comparison with the overlaps generated under a random distribution of such regions in the analyzed double round-robin (DRR) populations HvDRR13, HvDRR27, and HvDRR28. The random distribution of the genomic regions was simulated with 1,000 permutations.

| Recombination region | Genetic region | Population | Observed overlaps | Random overlaps |
| --- | --- | --- | --- | --- |
| Coldspot windows | Genes | HvDRR13 | 23989 | 37029 |
|  |  | HvDRR27 | 19545 | 26369 |
|  |  | HvDRR28 | 24832 | 37091 |
|  | Intergenic regions | HvDRR13 | 77006 | 129894 |
|  |  | HvDRR27 | 73553 | 87290 |
|  |  | HvDRR28 | 78332 | 135060 |
| Hotspot windows | Genes | HvDRR13 | 1057*** | 237 |
|  |  | HvDRR27 | 739*** | 146 |
|  |  | HvDRR28 | 1264*** | 282 |
|  | Intergenic regions | HvDRR13 | 823 | 840 |
|  |  | HvDRR27 | 684*** | 486 |
|  |  | HvDRR28 | 1057 | 1031 |

\*\*\* $P < 0.001$ for $H_0: \mu_{Observed} \leq \mu_{Random}$ ; Z-test.

### The variation in methylation and SVs in coldspot and hotspot windows among barley populations

Significant differences (*P <* 0.016) were observed in the methylation levels of the three analyzed populations, either by analyzing the methylated sequence contexts separately or by analyzing their average (Figure 3). Such differences among populations observed for the total windows were also detected in the coldspot windows in all of the analyzed chromosome regions. In contrast, the methylation level in the hotspot windows was found to be equal (*P >* 0.016) among populations in any of the analyzed chromosome regions, either by analyzing the methylated sequence contexts separately or their average. A similar trend was observed for the total SV load fraction: while observing significant differences (*P <* 0.016) among populations in the total windows but also in the coldspots of both the pericentromeric region and the distal subregions, such differences were not observed for the hotspot windows.

The methylation level of the two parental inbred lines of each of the analyzed populations differed significantly (*P <* 0.008) at the CpG and CHG sequence contexts of the genomic windows identified as coldspots in their respective offspring (Table 2). Thus, the increased methylation of only one of the parental genotypes at a genomic region might be enough to generate a coldspot in the offspring. In hotspot windows, no significant differences (*P >* 0.008) between parental inbred lines were found in the methylation level at any of the three analyzed sequence contexts, indicating that parents must have equally low methylation at a genomic segment to allow a recombination hotspot.

**Table 2:** Comparison of the methylation level at sequence contexts CpG, CHG, and CHH in the windows identified as recombination hotspots and coldspots in the double round-robin (DRR) populations HvDRR13, HvDRR27, and HvDRR28 for the respective parental inbred lines. The coldspot and hotspots windows were calculated on the basis of the recombination rate of the DRR populations. For a given population and given methylated sequence context, the significant difference (α = 0.016) in the Wilcoxon’s rank sum test among genomic window groups are indicated with different letters.

| Population | Coldspot windows |  |  |  | Hotspot windows |  |  |  |
| --- | --- | --- | --- | --- | --- | --- | --- | --- |
|  | Offspring | HOR8160 | Unumli-Arpa | SprattArcher | Offspring | HOR8160 | Unumli-Arpa | SprattArcher |
| HvDRR13 |  |  |  |  |  |  |  |  |
| CpG | 0.880 c | 0.875 b | - | 0.883 a | 0.666 a | 0.661 a | - | 0.668 a |
| CHG | 0.581 c | 0.571 b | - | 0.590 a | 0.334 a | 0.322 a | - | 0.345 a |
| CHH | 0.020 c | 0.020 b | - | 0.021 a | 0.029 a | 0.028 a | - | 0.030 a |
| HvDRR27 |  |  |  |  |  |  |  |  |
| CpG | 0.894 b | - | 0.891 a | 0.892 b | 0.673 a | - | 0.667 a | 0.678 a |
| CHG | 0.595 c | - | 0.587 a | 0.602 b | 0.346 a | - | 0.330 a | 0.362 a |
| CHH | 0.019 b | - | 0.019 a | 0.020 b | 0.029 ab | - | 0.027 a | 0.031 b |
| HvDRR28 |  |  |  |  |  |  |  |  |
| CpG | 0.880 c | 0.875 b | 0.881 a | - | 0.645 a | 0.643 a | 0.644 a | - |
| CHG | 0.573 c | 0.570 b | 0.574 a | - | 0.319 a | 0.319 a | 0.315 a | - |
| CHH | 0.019 a | 0.019 b | 0.019 a | - | 0.028 a | 0.028 a | 0.027 a | - |

## DISCUSSION

### Detection of recombination events at high-resolution in barley

The substantial decrease of the median inter-SNP distance compared to a previous study with the same three populations (Casale et al., 2022) produced a slight increase of the CO discovery rate of 0.31 times when considering the COs related to *>* 3 Mbp marker blocks. However, such increase jumps to 2.74 and 10.59 times, if considering the COs related to *>* 500 kb and *>* 10 kb blocks, respectively (Tables S3 and S5). The assumed part of the observed increase due to the additional recombination that occurred at heterozygous regions in the selfing generation analyzed in the first study is expected to be small due to the decreasing remaining heterozygosity after every selfing generation that produces fewer new observable COs per generation during inbreeding (Esch et al. 2007). By analyzing the detected CO rate between comparable low and high-resolution analyses reported in previous studies, only small differences were detected in populations of *Arabidopsis thaliana* and maize (*Zea mayz*) (McMullen et al. 2009; Rodgers-Melnick et al. 2015), but substantial differences were reported in populations of wheat (Esch et al. 2007; Gutierrez-Gonzalez et al. 2019; Gardiner et al. 2019) and *Populus* (Apuli et al. 2020). In addition to the different utilized resolutions, other reasons behind observing differences in the recombination rate in the same population may include the genotyping error rate, the data filtering criteria, and the size of considered CO events.

The observed CO rate per RIL per chromosome per generation in the present study, when considering the COs related to *>* 3 Mbp marker blocks, is in line with high-resolution studies in *Arabidopsis* (Sun et al. 2012; Lu et al. 2012; Yang et al. 2012; Wijnker et al. 2013; Qi et al. 2014; Rowan et al. 2019) and rice (*Oryza sativa*)(Si et al. 2015) that reported rates of 1.5–2.2 COs per chromosome per generation, and with another high-resolution study on wheat (Gardiner et al. 2019) when looking at the COs related to *>* 500 kb marker blocks.

The COs per RIL per chromosome per generation observed in our study when considering all COs related to blocks (10 kb) should be compared with values observed by Yang et al. (2012) in *Arabidopsis* and Gardiner et al. (2019) in wheat. In such comparisons, however, considering every marker block shorter than 10 kb as a GC is an arbitrary threshold. This has the potential to cause misclassification between COs and GCs among the categories of COs related to blocks *>* 10 kb and those related to GCs between 2 and 10 kb.

The present study is the first to characterize GC events in barley, along with a few in other crop species (Li et al. 2015; Si et al. 2015; Gardiner et al. 2019). The poor documentation of GCs in plants beyond studies in *Arabidopsis* is because phenotypic screens can barely detect GC events. In addition, the detection of GC events at the molecular genetic level is also challenging because of their short length (Mancera et al. 2009; Mercier et al. 2015). This makes GC detection very sensitive to the marker density and GC rate, which also depend on the recombination rate, the tract length of the repair intermediates where GCs occur, and the polymorphism density (Wijnker et al. 2013). Moreover, in most flowering plants, gametes do not remain grouped after meiosis, making it difficult to observe the expected 3:1 allelic proportion of GCs (Sun et al. 2012). In the present study, the average number of genome-wide GC events per RIL across populations was 58, 251, and 6,521 for long (2–10 kb), medium (20 bp–2 kb), and short (2–20 bp) GC-related marker block lengths, respectively. The SNPs analyzed per population could be translated to 0.00003, 0.00014, and 0.0039 GC events per site per RIL per generation for the different types of GC-related marker block lengths. Marker blocks shorter than 20 bp are expected to contain a high number of false-positive GCs since they were predominantly called based on two markers only. Therefore, if considering the GC-related marker blocks of long (2–10 kb) and medium (20 bp–2 kb) length only, the detected 0.00017 GCs per site per RIL per generation is on the same order of magnitude as that observed in studies using a similar approach for GC detection (Yang et al. 2012; Gardiner et al. 2019). However, the observed GC rate in our study was two orders of magnitude greater than that reported in tetrad analysis studies performed in *Arabidopsis* (Lu et al. 2012; Sun et al. 2012; Wijnker et al. 2013) and sequencing of rice F2 populations (Si et al. 2015). This disagreement might be explained not only by the less precise GC detection method used in our study but also by the occurrence of nonallelic sequence alignments caused by SVs inflating the number of false-positive gene conversions (Qi et al. 2014; Si et al. 2015). It is also worth noting that the reported GC rate is the frequency of GCs generated from NCO and CO events combined. In the present study, we did not attempt to estimate the rate of NCO in barley because these NCOs are only traceable after gamete formation when they lead to GC, and it was not possible to precisely differentiate between CO and NCO conversion tracts with the employed marker resolution. In addition, the DSB rate in barley is not known enough to estimate the NCO rate from the detected CO events.

The present study is the first to report genome-wide recombination hotspots at high-resolution in barley. On average, across the three investigated populations, of the 80 hotspots per chromosome, only 12 were found in the pericentromeric region, and the rest were found in the distal regions of the barley chromosomes (Figure S10 and Table S8). In addition, while the three investigated populations always shared one parent, the proportion of shared hotspots between two and three populations was 10% and 1%, respectively, of the total hotspots detected in a given population (Figure S11A). This observation is in good agreement with previous studies in *Arabidopsis* showing that recombination hotspots were cross-specific (Salomé et al. 2012).

In contrast, in the case of the GC hotspot windows, the overlap among the three populations was more than 15% (Figure S12). Moreover, GC hotspots were found to have high overlap not only with GC hotspots of the same population but also with windows that are hotspots in other populations (Table S10). Thus, GC hotspots might be considered fingerprints of population-specific silenced COs that result in NCO in regions with high DSB rates in the genome of the species. This observation suggested that although the CO rate and distribution present extensive intraspecies variation, such DSBs might be highly conserved within the species. However, this requires further research.

Additionally, in barley, recombination hotspots alternate in the genome with coldspots. For example, in domains where CO rates are significantly lower than the genome average (Figure S10), as observed in previous studies in other species (Mercier et al. 2015). Indeed, by dividing the genome into 10 kb genomic windows, hotspot windows were found to be adjacent to coldspot windows in 42.5% of the cases in the distal regions of the barley chromosomes (Table S15). This continuos intermittence in the recombination rate might explain the above-mentioned large difference in CO events found between the high- and low-resolution analyses on these barley populations.

To avoid calling recombination coldspots in the pericentromeric and telomeric regions of the chromosomes, which are long regions depleted from recombination as seen in previous studies (Boideau et al. 2022), in the present study, coldspots were identified only in the distal region of the chromosomes by employing a long physical distance margin between regions. The majority of the detected 10 kb coldspot windows were located in coldspot regions with an average length of 322 kb (Table S7). Each population shared 60% of its coldspot windows and more than 16% with the other two populations, thus demonstrating a greater conservation of coldspots than hotspots in barley (Figure S11B). Such differential conservation between recombination hotspots and coldspots might be linked to the different genomic features determining their occurrence.

### The genomic features associated with recombination rate variation in barley

The present study is the first comprehensive evaluation of the genomic features associated with differences in recombination rates among barley populations. On a scale of 1 Mbp windows, the recombination rate was found to be positively correlated with sequence divergence among parental inbred lines, gene density, and SV load on the barley chromosomes and was negatively correlated with the methylation level (Figure 1).

The results of the present study are in line with earlier studies in plants in which recombination was found to be positively associated with genetic divergence among homologous chromosomes (Yang et al. 2012; Marand et al. 2019; Blackwell et al. 2020). Although a negative association was reported in some studies (Saintenac et al. 2011; Gion et al. 2016; Bouchet et al. 2017; Serra et al. 2018; Jordan et al. 2018; Gutierrez-Gonzalez et al. 2019), the contradiction might be explained by a sigmoid relationship between both variables, meaning that recombinatiom has a positive correlation with genetic divergence until a level after which the high polymorphism among homologs suppresses COs due to the increase in mismatches, as recently reported in *Arabidopsis* (Blackwell et al. 2020; Hsu et al. 2022). In this respect, the few observed 1 Mbp windows with a significant negative association were found to have parental sequence divergence at the relative maximum level, which appeared to be associated with an extensive SV load (Figure S5A and B).

The SV load was not identified as a determining factor for the differences in the recombination rate among populations, presumably because the employed resolution of 1 Mbp was too coarse to detect differences among populations, as most of the analyzed SVs were smaller (Figure 2). Additionally, the positive correlation between the recombination rate and the SV load on a broad scale might be explained by the accumulation of DNA repair errors in highly recombining regions throughout the evolutionary history of barley (Figures 1A and 1C). Genomic regions with a high rate of DSBs are expected to have a historically increased mutation rate produced by COs and GCs, which elevates the allelic diversity at such regions among genotypes as demonstrated in humans (Arbeithuber et al. 2015; Halldorsson et al. 2019).

In addition to the positive correlation shown between recombination and gene density on a broad scale, in the present study, 10 kb hotspot windows were found to be associated with regions of high gene density (Table 1), as repeatedly reported in previous studies in grasses (Rodgers-Melnick et al. 2015; Darrier et al. 2017; Bouchet et al. 2017; Jordan et al. 2018; Gardiner et al. 2019; Marand et al. 2019; Casale et al. 2022), and other plant families (Paape et al. 2012; Choi et al. 2013; Silva-Junior and Grattapaglia 2015; Gion et al. 2016; Wang et al. 2016; Apuli et al. 2020). Furthermore, the hotspot windows were located in proximity (*<* 20 kb apart) but did not overlap with the genes (Figure 4). This finding in barley is in line with previous observations in *Arabidopsis*, maize, and rice showing an increased CO frequency toward gene promoters and terminators (Choi et al. 2013; Wijnker et al. 2013; Li et al. 2015; Marand et al. 2019; Sun et al. 2019), similar to that observed in budding yeast (*Saccharomyces cerevisiae*) (Pan et al. 2011).

In the present study, the genetic effects were calculated as the proportion of the sum of the *GRE* of both parents for a given population that was not explained by methylation, assuming parental sequence divergence and SV load as part of the *SRE* (Casale et al. 2022). Such genetic effects were shown to be the factor explaining the greatest proportion of differences in recombination rates among barley populations. Here, we hypothesize that such genetic effects are the product of the expression of genes related to the recombination machinery being in part modulated by the methylation level, explaining a portion of the uneven distribution of the recombination hotspots along the barley chromosomes. The wide distribution of the observed genetic effects along the chromosomes is in line with previous studies in wheat (Jordan et al. 2018) and barley (Casale et al. 2022) reporting that differences in the genome recombination rate among populations are explained by multiple loci with small effects.

A negative correlation was observed on a broad scale between recombination rate and the extent of methylation in the CpG and CHG sequence contexts (Figure 1B). This is in accordance with previous studies in other plant species showing that COs occurred in euchromatic regions while heterochromatic regions were depleted of COs and that hypomethylation at CpG sites increased the genome-wide CO rate (Melamed-Bessudo and Levy 2012; Colomé-Tatché et al. 2012; Salomé et al. 2012; Yelina et al. 2012; Mirouze et al. 2012; Wijnker et al. 2013; Rodgers-Melnick et al. 2015; Marand et al. 2019; Apuli et al. 2020; Boideau et al. 2022; Fernandes et al. 2024). This association was confirmed at high-resolution by oberving the relationship between 10 kb coldspot windows and increased methylation in the CpG and CHG sequence contexts (Figure 3). Furthermore, compared with those in the nonhotspot windows, the methylation levels in the CpG and CHG sequence contexts in both the distal and pericentromeric regions of the barley chromosomes decreased in the hotspot windows. This result is in line with previous findings in maize showing a strong relationship between the occurrence of hotspots and decreased CpG and CHG methylation but no association with CHH methylation (Rodgers-Melnick et al. 2015). In this respect, it was suggested that increased recombination was associated with increased CHH methylation in regions with high CpG-related methylation levels but with decreased recombination where CpG methylation was low (Rodgers-Melnick et al. 2015).

The presence of SVs was shown to suppress COs in previous studies in *Arabidopsis* (Rowan et al. 2019; Fernandes et al. 2024) and other plant species (Rodgers-Melnick et al. 2015; Shen et al. 2019). In the present study, we were able to detect a decreased overlap and a longer distance between CO breakpoints and SVs compared to a random distribution, thus indicating the negative association of SVs with the occurrence of COs in barley (Tables S13 and S14). The type and size of the SVs were not found to be related to such effects, which is in line with previous findings in *Arabidopsis* (Rowan et al. 2019). Moreover, this is the first study showing the joint effect of methylation and the accumulation of SVs in determining genomic regions deprived of recombination outside the pericentromeric region (Fernandes et al. 2024) and the variation in this effect within the genomic region (Figure 3 and Tables S11 and S12). In the distal telomeric region of the barley chromosomes, most 10 kb coldspot windows were found to be associated with increases in both the recombination level and the SV load. However, in the distal proximal region, increased methylation was found to be associated with most of the 10 kb coldspot windows, but an increased SV load was found to be associated with coldspot windows with no increased methylation. Interestingly, the effects of both methylation and SVs on the occurrence of coldspots were noticeable not only when comparing coldspot windows with other windows located far away in the same genomic region but also when comparing coldspots with their neighboring windows. This indicated a marked local effect of methylation and SVs on recombination suppression (Figure 4). In addition, the differences in both methylation level and SV load among barley populations were found to be responsible for the differences in the localization of coldspot windows among such populations. The parental inbred lines of the analyzed populations were found to differ in the methylation level in the genomic windows identified as coldspots in their offspring populations (Table 2), indicating that the inheritance of high methylation from only one parent was sufficient to prevent recombination in a particular region of the genome.

In a previous study on the same barley populations in which methylation was not separated from the genetic effects of genotypes, the effect of individual parental inbred lines was shown to be the major determinant of the recombination rate of the respective biparental offspring populations (Casale et al. 2022). The increased methylation at genomic regions leading to coldspots might be an important part of the genetic effect of the parents on the recombination rate, which was negatively correlated with methylation in the present study.

In contrast to the above described association of methylation and SVs with the occurence of recombination coldspots, no significant differences between parental inbred lines were found in the methylation level of hotspot windows, indicating that parents must have equally low methylation at a genomic segment to allow a recombination hotspot. The employed window resolution of 10 kb, which is longer than the length reported earlier for recombination hotspots in other plant species (Choi and Henderson 2015), might be the reason for the lack of detection of SV effects on hotspots and the lack of differences in methylation between hotspots and their neighboring windows.

Our findings demonstrate that the recombination landscape in barley is highly predictable. Most of the recombination occurs in multiple short highly recombining sections in the distal regions of the chromosomes. These recombination hotspots are located in proximity to genes and where the levels of methylation and SV load are low enough to allow CO concretion. In this sense, such hotspots alternate with long regions deprived of recombination because of increased methylation or the accumulation of SVs preventing CO from occurring. Therefore, local differences in the recombination rate among barley populations can be explained to a considerable extent by differences in the methylation level and the accumulation of SVs at multiple locations within the genome. Such differences are highly inheritable and can be determined by the effect of only one parent in a cross. However, our analyses suggest that in addition to these two genomic features, additional differences in the recombination machinery must exist, which forms the basis for what we designated genetic effect.

## METHODS

### Identification of the genomic features associated with recombination rate variation among barley populations

The recombination rates of 45 biparental barley populations, referred to as double round-robin populations, were obtained from Casale et al. (2022). These have been derived from genotyping the populations with the 50K Illumina Infinium iSelect SNP genotyping array (Bayer et al. 2017). The recombination rates were recalculated based on the Morex v3 reference genome sequence (Mascher et al. 2021) at 1 Mbp genomic windows. The pericentromeric region of each chromosome was defined as the continuous region surrounding the centromere for which the average recombination rate across the 45 DRR populations was 5-fold lower than the respective chromosome average across populations in 1 Mbp genomic windows. The regions of the chromosome that did not belong to the pericentromeric region were designated in the following as distal regions.

Whole-genome bisulfite DNA sequencing data for the 23 DRR parental inbred lines was obtained by extracting DNA from inbred lines from a mix of tissues, including the whole seedling plant, the leaf, and the apex, at stage 47 on the Zadoks scale. DNA library preparation was performed with NEBNext® Ultra™ II (New England Biolabs, Inc., USA), and bisulfite conversion was performed with the EZ DNA Methylation-Lightning Kit (Zymo Research, USA). The resulting 150 bp paired-end libraries were sequenced with Illumina HiSeq™ 2000 and NovaSeq™ (Il-lumina, Inc., USA). The raw reads were trimmed with Trim Galore! (Krueger et al. 2023), mapped against the Morex v3 reference genome with Bismark (Krueger and Andrews 2011), and aligned with Bowtie 2 (Langmead and Salzberg 2012). For quality control, SNPs were called with Bis-SNP (Liu et al. 2012) and compared with single nucleotide variation (SNV) data generated by DNA sequencing of the respective inbred lines (Weisweiler et al. 2022). The level of methylation in cytosine positions present in the methylated sequence contexts CpG, CHG, and CHH, was calculated as the percentage of the methylated reads per position. For each DRR population, the methylation level at each sequence context in a given genomic window was calculated as the average among the respective parental inbred lines’ methylation level values for the methylated cytosine positions in that window, weighted by the number of methylated cytosine positions corresponding to each parent. The average methylation level across the three sequence contexts in a given genomic 1 Mbp window was calculated as the average among the calculated methylation levels for such contexts in the window, weighted by the number of methylated cytosine positions corresponding to each of the contexts in the window. The difference in methylation level between the two parental inbred lines of a population at each 1 Mbp genomic window was calculated for the three methylated sequence contexts and their average.

The gene density in 1 Mbp windows was calculated as the physical fraction spanned by the coding sequence of genes in each window. The locations of genes and intergenic regions on the barley chromosomes were obtained from the Morex v3 reference sequence. The genetic divergence among parents of the DRR populations was calculated from single nucleotide variant (SNV) data derived from genome-wide sequencing (Weisweiler et al. 2022).

The SNVs were also used to calculate the general recombination effects (*GRE*) of the parental inbred lines as described by Casale et al. (2022). In the next step, the proportion of the sum of the GRE of both parents for a given population that was not explained by the average methylation in each 1 Mbp window was estimated using linear regression. This residual was assumed to represent the genetic effects on recombination in a given genomic window. The specific recombination effect (*SRE*) for a given parental combination was not taken into account to estimate genetic effects because it was previously described to cause only a minor effect on the recombination rate of a given biparental barley population (Casale et al. 2022). The SVs such as inversions, insertions, deletions, duplications, and translocations, between the parental inbred lines of the DRR populations and the Morex reference genome were obtained from Weisweiler et al. (2022). The SVs were categorized by size (50—299 bp, 0.3—4.9 kb, 5—49 kb, 50—249 kb, 0.25—1 Mbp, and *>*1 Mbp), except for translocations whose length was not determined (Table S1). The physical length fraction spanned by SVs in every 1 Mbp genomic window across the genome was estimated for all SV categories and sizes. Furthermore, the total SV span fraction generated by the sum of all SV categories and sizes in each 1 Mbp window was calculated (hereafter referred to as SV load).

Pearson’s correlation between all pairs of the abovementioned genomic features was calculated by genomic window and population. To identify which of the features better explained the recombination rate variation among the 45 DRR populations at each 1 Mbp genomic window in the barley chromosomes, a stepwise regression approach was used. The procedure keeps for each 1 Mbp window the subset of genomic features that explain differences among the recombination rates of the DRR populations with the highest Akaike information criterion (AIC). The fraction of 1 Mbp windows of the barley genome in which a given genomic feature was retained in the model provides an estimation of the feature’s importance across the entire genome. Moreover, the direction of the correlation between the analyzed features and the recombination rate provides a notion of the impact of the feature on either promoting or repressing recombination. The model included the total SV load, parental sequence divergence, genetic effects, average methylation level, and difference among parental inbred lines at the average methylation level. In addition, an extended model was constructed by breaking down the methylation-related variables into the respective methylated sequence contexts and the SV load into the different SV types.

### Investigation of the genomic features associated with the recombination rate in barley at high-resolution Plant material, genotyping, and data cleaning

From the 45 DRR population set, three populations (HvDRR13, HvDRR27, and HvDRR28) were selected for high-density genotyping using an mRNAseq approach as described by Arlt et al. (2023). These populations are the product of a triangle cross among three parental inbred lines (HOR8160, SprattArcher, and Unumli-Arpa). The respective 64, 92, and 79 RILs were cultivated at the S7 generation in petri dishes in a randomized incomplete block design where the parental inbred lines were included as controls. The blocks were harvested 7 days after planting with less than 2 hours difference between the first and last sample. The whole seedling was utilized for mRNA extraction. For mRNA sequencing (RNA-Seq), the RIL-specific library was constructed using the VAHTS Universal V6 RNA-seq Library Prep Kit for Illumina (Vazyme, China), and RIL-specific barcodes were used. The pooled libraries were sequenced on the DNBSEQ-G400 platform (MGI Tech Co., Ltd., China) by BGI Genomics (Beijing, China), generating 1.42 billion 150 bp paired-end reads. The reads were trimmed with Trimmomatic (Bolger et al. 2014) and aligned to the Morex v3 reference sequence using HISAT2 (Kim et al. 2019). Variant calling was performed using BCFtools (Li et al. 2009). The obtained variants were selected based on their intersection with the reported SNVs from the parental inbred lines (Weisweiler et al. 2022). Furthermore, a 12 K subset of the SNP markers reported previously (Casale et al. 2022) for the same parental inbred lines and RILs was added to the total set at genomic positions not present in the RNAseq dataset. On a population basis, SNPs carrying nonparental alleles were set to missing data, and SNPs with 100% missing data or monomorphic parental alleles were discarded. Missing data for genotypes at polymorphic positions were reconstructed using Beagle (Browning et al. 2018) with default parameters.

### Detection of recombination events

A recombination event in a given RIL haplotype was called when a block of SNP alleles inherited from one parental inbred switched to a block of SNP alleles belonging to the other parent (i.e., parental allele phase change). The recombination breakpoints were determined as the midpoint of the region between both blocks (i.e., the CO interval). The blocks comprising fewer than three SNPs were considered false positive CO events and were discarded. Then, blocks shorter than 10 kb were considered GC events, while blocks longer than 10 kb were considered to be produced by CO (Yang et al. 2012; Gardiner et al. 2019). To enable comparisons with earlier studies (Yang et al. e.g. 2012; Gardiner et al. e.g. 2019), GC-related blocks were grouped into long (2–10 kb), short (20 bp–2 kb), and very short (2–19 bp) GC blocks, while the CO-related segments were grouped into short (10–500 kb), medium (500 kb–3 Mbp), and long (*>* 3 Mbp) CO blocks. The longest block length threshold that kept every major parental allele phase change (3 Mbp) was defined visually by graphical genotypes on a 500 kb scale from 0.5 to 20 Mbp (Figure S2). The CO block length categories were considered different CO layers, and further analyses were performed on a multilayer basis. Individuals with a CO count falling outside the 3-fold interquartile range of their respective population were assumed to be outliers and were discarded from further analyses.

The pericentromeric and distal regions of each chromosome were defined as explained above at 10 kb genomic windows for each of the three analyzed populations independently. The distal regions were defined as the chromosome segments between the pericentromeric region and the telomeres of the chromosomes.

In each 10 kb genomic window of the genome, the physical fraction spanned by the CO intervals determined in all RILs of a population was aggregated to calculate the accumulated CO probability per window on a population basis. The accumulated CO probabilities per window were normalized per chromosome and per population. The CO hotspot windows in a given population were defined as the windows with a CO probability *>* 99%. The GC hotspot windows were determined in the same way as the CO hotspot windows. The windows located in the distal regions with a CO probability of zero were considered coldspot windows. The coldspot windows that were located beside other coldspot windows were combined into coldspot regions. To avoid calling coldspot windows near the pericentromeric region and telomeres, only windows located away from such regions were considered coldspot windows. This distance was granted by introducing an arbitrary margin with a length of 12.5% of the physical length of the respective distal region. To control for spurious associations generated by the variation of the features along the chromosomes, the windows from the distal regions were grouped into those close to the telomere (distal telomeric) and those close to the pericentromeric region (distal proximal).

### Investigation of the association between genomic features and recombination rate

The fraction of each window spanned by SVs, the gene density, the methylation level at the sequence contexts CpG, CHG, and CHH, their average, and the parental difference for these contexts were calculated as described above for each 10 kb genomic window of the barley chromosomes in the three analyzed populations HvDRR13, HvDRR27, and HvDRR28. In addition, the 10 kb windows neighboring coldspot and hotspot regions (any genomic length spanned by contiguous coldspot or hotspot 10 kb windows) were grouped by their relative position in the range from −40 kb to +40 kb around the respective coldspot or hotspot in the mentioned chromosome regions. The statistical comparison among window groups of any kind for the mentioned features was performed with the Mann–Whitney *U* test with Bonferroni correction for multiple testing.

The observed overlaps of the coldspot and hotspot regions with genes and intergenic regions in the chromosomes of the three analyzed populations were statistically compared with random overlaps simulated with regioneR (Gel et al. 2016) by running 1, 000 permutations on a by-chromosome basis where the respective pericentromeric regions were masked.

The observed overlap between CO intervals and the genomic regions spanned by SVs was assessed as described above with regione R for insertions/deletions, duplications, and inversions. In addition, the mean distance between the CO breakpoints and their closest SV in the genome was compared with the equivalent of random simulated COs in 10 Mbp genomic windows, each with a probability of CO occurrence according to the recombination rate per chromosome in the DRR populations reported by Casale et al. (2022). The significant differences among the means of the observed and simulated CO-SV distances were evaluated with the Mann–Whitney *U* test.

## DATA ACCESS

The methylation data of the 23 DRR population parental lines and the raw mRNA sequencing data of the RILs from populations HvDRR13, HvDRR27, and HvDRR28 (and their parental lines) are both available at NCBI, BioProject accessions PRJNA1100572 and 1088431, respectively. Employed scripts are available from the authors upon request.

## ACKNOWLEDGEMENTS

Computational infrastructure and support were provided by the Centre for Information and Media Technology at Heinrich Heine University Düsseldorf. The authors give thanks to the IPK for providing the seeds of the diversity panel. We thank our former colleagues Andrea Lossow, Nicole Kliche-Kamphaus, Nele Kaul, Isabelle Scheibert, Marianne Haperscheid, George Alskief, Florian Esser as well as our present colleagues Konstantin Shek and Stefanie Krey for their excellent technical assistance in creating and maintaining the DRR populations.

## AUTHORS’ CONTRIBUTIONS

FC contributed to the design of the project, analysed the data and performed the analyses related to meiotic recombination, and performed all statistical analyses. CA created the libraries, sequenced, and processed the RNA-Seq data. MK analysed the bisulfite-sequenced DNA data. Both CA and MK contributes equally to this work as second authors. JL created the segregating populations. JE and TH created the bisulfite libraries for DNA sequencing. BS designed and coordinated the project. FC and BS wrote the manuscript. All authors read and approved the final manuscript.

## REFERENCES

Apuli RP, Bernhardsson C, Schiffthaler B, Robinson KM, Jansson S, Street NR, and Ingvarsson PK. 2020. Inferring the genomic landscape of recombination rate variation in european aspen (Populus tremula). G3: Genes, Genomes, Genetics 10: 299–309.

Arbeithuber B, Betancourt AJ, Ebner T, and Tiemann-Boege I. 2015. Crossovers are associated with mutation and biased gene conversion at recombination hotspots. Proceedings of the National Academy of Sciences 112: 2109–2114.

Arlt C, Wachtmeister T, Köhrer K, and Stich B. 2023. Affordable, accurate and unbiased RNA sequencing by manual library miniaturization: A case study in barley. Plant Biotechnology Journal 21: 2241–2253.

Barton NH and Charlesworth B. 1998. Why sex and recombination? Science 281: 1986–1990.

Baudat F and Massy BD. 2007. Regulating double-stranded dna break repair towards crossover or non-crossover during mammalian meiosis. Chromosome research 15: 565–577.

Bauer E, Falque M, Walter H, Bauland C, Camisan C, Campo L, Meyer N, Ranc N, Rincent R, Schipprack W, et al. 2013. Intraspecific variation of recombination rate in maize. Genome Biology 14.

Bayer MM, Rapazote-Flores P, Ganal M, Hedley PE, Macaulay M, Plieske J, Ramsay L, Russell J, Shaw PD, Thomas W, et al. 2017. Development and evaluation of a barley 50k iselect SNP array. Frontiers in Plant Science 8: 1792.

Blackwell AR, Dluzewska J, Szymanska-Lejman M, Desjardins S, Tock AJ, Kbiri N, Lambing C, Lawrence EJ, Bieluszewski T, Rowan B, et al. 2020. MSH2 shapes the meiotic crossover landscape in relation to interhomolog polymorphism in Arabidopsis. The EMBO Journal 39: 1–22.

Blary A and Jenczewski E. 2019. Manipulation of crossover frequency and distribution for plant breeding. Theoretical and Applied Genetics 132: 575–592.

Boideau F, Richard G, Coriton O, Huteau V, Belser C, Deniot G, Eber F, Falentin C, de Carvalho JF, Gilet M, et al. 2022. Epigenomic and structural events preclude recombination in Brassica napus. New Phytologist 234: 545–559.

Bolger AM, Lohse M, and Usadel B. 2014. Trimmomatic: A flexible trimmer for illumina sequence data. Bioinformatics 30: 2114–2120.

Bouchet S, Olatoye MO, Marla SR, Perumal R, Tesso T, Yu J, Tuinstra M, and Morris GP. 2017. Increased Power To Dissect Adaptive Traits in Global Sorghum Diversity Using a Nested Association Mapping Population. Genetics 206: 573– 585.

Browning BL, Zhou Y, and Browning SR. 2018. A one-penny imputed genome from next-generation reference panels. American Journal of Human Genetics 103: 338–348.

Burt A. 2000. Perspective: Sex, recombination, and the efficacy of selection - was weismann right? Evolution 54: 337–351.

Casale F, Van Inghelandt D, Weisweiler M, Li J, and Stich B. 2022. Genomic prediction of the recombination rate variation in barley – A route to highly recombinogenic genotypes. Plant Biotechnology Journal 20: 676–690.

Choi K and Henderson IR. 2015. Meiotic recombination hotspots - a comparative view. Plant Journal 83: 52–61.

Choi K, Zhao X, Kelly KA, Venn O, Higgins JD, Yelina NE, Hardcastle TJ, Ziolkowski PA, Copenhaver GP, Franklin FCH, et al. 2013. Arabidopsis meiotic crossover hot spots overlap with H2A.Z nucleosomes at gene promoters. Nature Genetics 45: 1327–1338.

Colomé-Tatché M, Cortijo S, Wardenaar R, Morgado L, Lahouz B, Sarazin A, Etcheverry M, Martin A, Feng S, Duvernois-Berthet E, et al. 2012. Features of the arabidopsis recombination landscape resulting from the combined loss of sequence variation and dna methylation. Proceedings of the National Academy of Sciences of the United States of America 109: 16240–16245.

Darrier B, Rimbert H, Balfourier F, Pingault L, Josselin AA, Servin B, Navarro J, Choulet F, Paux E, and Sourdille P. 2017. High-resolution mapping of crossover events in the hexaploid wheat genome suggests a universal recombination mechanism. Genetics 206: 1373–1388.

Dreissig S, Mascher M, Heckmann S, and Purugganan M. 2019. Variation in recombination rate is shaped by domestication and environmental conditions in barley. Molecular Biology and Evolution 36: 2029–2039.

Dreissig S, Maurer A, Sharma R, Milne L, Flavell AJ, Schmutzer T, and Pillen K. 2020. Natural variation in meiotic recombination rate shapes introgression patterns in intraspecific hybrids between wild and domesticated barley. New Phytologist 228: 1852–1863.

Esch E, Szymaniak JM, Yates H, Pawlowski WP, and Buckler ES. 2007. Using crossover breakpoints in recombinant inbred lines to identify quantitative trait loci controlling the global recombination frequency. Genetics 177: 1851–1858.

Fernandes JB, Naish M, Lian Q, Burns R, Tock AJ, Rabanal FA, Wlodzimierz P, Habring A, Nicholas RE, Weigel D, et al. 2024. Structural variation and dna methylation shape the centromere-proximal meiotic crossover landscape in arabidopsis. Genome Biology 25: 30.

Gardiner LJ, Wingen LU, Bailey P, Joynson R, Brabbs T, Wright J, Higgins JD, Hall N, Griffiths S, Clavijo BJ, et al. 2019. Analysis of the recombination landscape of hexaploid bread wheat reveals genes controlling recombination and gene conversion frequency. Genome Biology 20: 69.

Gel B, Díez-Villanueva A, Serra E, Buschbeck M, Peinado MA, and Malinverni R. 2016. RegioneR: An R/Bioconductor package for the association analysis of genomic regions based on permutation tests. Bioinformatics 32: 289–291.

Gion JM, Hudson CJ, Lesur I, Vaillancourt RE, Potts BM, and Freeman JS. 2016. Genome-wide variation in recombination rate in Eucalyptus. BMC Genomics 17: 1–12.

Giraut L, Falque M, Drouaud J, Pereira L, Martin OC, and Mézard C. 2011. Genome-wide crossover distribution in arabidopsis thaliana meiosis reveals sex-specific patterns along chromosomes. PLoS Genetics 7: e1002354.

Gutierrez-Gonzalez JJ, Mascher M, Poland J, and Muehlbauer GJ. 2019. Dense genotyping-by-sequencing linkage maps of two Synthetic W7984×Opata reference populations provide insights into wheat structural diversity. Scientific Reports 9: 1–15.

Hall JC. 1972. Chromosome segregation influenced by two alleles of the meiotic mutant c(3)G in Drosophila melanogaster. Genetics 71: 367–400.

Halldorsson BV, Palsson G, Stefansson OA, Jonsson H, Hardarson MT, Eggertsson HP, Gunnarsson B, Oddsson A, Halldorsson GH, Zink F, et al. 2019. Characterizing mutagenic effects of recombination through a sequence-level genetic map. Science 363.

Henderson IR. 2012. Control of meiotic recombination frequency in plant genomes. Current Opinion in Plant Biology 15: 556–561.

Higgins JD, Perry RM, Barakate A, Ramsay L, Waugh R, Halpin C, Armstrong SJ, and Franklin FCH. 2012. Spatiotemporal asymmetry of the meiotic program underlies the predominantly distal distribution of meiotic crossovers in barley. Plant Cell 24: 4096–4109.

Hsu YM, Falque M, and Martin OC. 2022. Quantitative modelling of fine-scale variations in the Arabidopsis thaliana crossover landscape. Quantitative Plant Biology 3: e3.

Jordan KW, Wang S, He F, Chao S, Lun Y, Paux E, Sourdille P, Sherman J, Akhunova A, Blake NK, et al. 2018. The genetic architecture of genome-wide recombination rate variation in allopolyploid wheat revealed by nested association mapping. Plant Journal 95: 1039–1054.

Kim D, Paggi JM, Park C, Bennett C, and Salzberg SL. 2019. Graph-based genome alignment and genotyping with HISAT2 and HISAT-genotype. Nature Biotechnology 37: 907–915.

Kim S, Plagnol V, Hu TT, Toomajian C, Clark RM, Ossowski S, Ecker JR, Weigel D, and Nordborg M. 2007. Recombination and linkage disequilibrium in Arabidopsis thaliana. Nature Genetics 39: 1151–1155.

Krueger F and Andrews SR. 2011. Bismark: a flexible aligner and methylation caller for bisulfite-seq applications. Bioinformatics 27: 1571–1572.

Krueger F, James F, Ewels P, Afyounian E, Weinstein M, Schuster-Boeckler B, Hulselmans G, and sclamons. 2023. Trimgalore v0.6.10.

Langmead B and Salzberg S. 2012. Fast gapped-read alignment with Bowtie 2. Nature methods 9: 357–359.

Lawrence EJ, Griffin CH, and Henderson IR. 2017. Modification of meiotic recombination by natural variation in plants. Journal of Experimental Botany 68: 5471–5483.

Li H, Handsaker B, Wysoker A, Fennell T, Ruan J, Homer N, Marth G, Abecasis G, and Durbin R. 2009. The sequence alignment/map format and SAMtools. Bioinformatics 25: 2078–2079.

Li X, Li L, and Yan J. 2015. Dissecting meiotic recombination based on tetrad analysis by single-microspore sequencing in maize. Nature Communications 6.

Liu Y, Siegmund K, Laird P, and Berman B. 2012. Bis-SNP: combined DNA methylation and SNP calling for bisulfite-seq data. Genome biology 13: R61.

Lu P, Han X, Qi J, Yang J, Wijeratne AJ, Li T, and Ma H. 2012. Analysis of Arabidopsis genome-wide variations before and after meiosis and meiotic recombination by resequencing landsberg erecta and all four products of a single meiosis. Genome Research 22: 508–518.

Mancera E, Bourgon R, Brozzi A, Huber W, and Steinmetz M. 2009. High-resolution mapping of meiotic crossovers and noncrossovers in yeast. Nature 454: 479–485.

Marand AP, Zhao H, Zhang W, Zeng Z, Fang C, and Jianga J. 2019. Historical meiotic crossover hotspots fueled patterns of evolutionary divergence in rice. Plant Cell 31: 645–662.

Martini E, Diaz RL, Hunter N, and Keeney S. 2006. Crossover homeostasis in yeast meiosis. Cell 126: 285–295.

Mascher M, Wicker T, Jenkins J, Plott C, Lux T, Koh CS, Ens J, Gundlach H, Boston LB, Tulpová Z, et al. 2021. Long-read sequence assembly: A technical evaluation in barley. Plant Cell 33: 1888–1906.

McMullen MD, Kresovich S, Villeda HS, Bradbury P, Li H, Sun Q, Flint-Garcia S, Thornsberry J, Acharya C, Bottoms C, et al. 2009. Genetic properties of the maize nested association mapping population. Science 325: 737–740.

Melamed-Bessudo C and Levy AA. 2012. Deficiency in dna methylation increases meiotic crossover rates in euchromatic but not in heterochromatic regions in Arabidopsis. Proceedings of the National Academy of Sciences of the United States of America 109.

Mercier R, Mézard C, Jenczewski E, Macaisne N, and Grelon M. 2015. The molecular biology of meiosis in plants. Annual Review of Plant Biology 66: 297–327.

Mirouze M, Lieberman-Lazarovich M, Aversano R, Bucher E, Nicolet J, Reinders J, and Paszkowski J. 2012. Loss of DNA methylation affects the recombination landscape in Arabidopsis. Proceedings of the National Academy of Sciences of the United States of America 109: 5880–5885.

Muller HJ. 1932. Some genetic aspects of sex. The American Naturalist 66: 118–138.

Mézard C. 2006. Meiotic recombination hotspots in plants. Biochemical Society Transactions 34: 531–534.

Nachman MW. 2002. Variation in recombination rate across the genome: Evidence and implications. Current Opinion in Genetics and Development 12: 657–663.

Paape T, Zhou P, Branca A, Briskine R, Young N, and Tiffin P. 2012. Fine-scale population recombination rates, hotspots, and correlates of recombination in the Medicago truncatula genome. Genome Biology and Evolution 4: 726–737.

Pan J, Sasaki M, Kniewel R, Murakami H, Blitzblau HG, Tischfield SE, Zhu X, Neale MJ, Jasin M, Socci ND, et al. 2011. A hierarchical combination of factors shapes the genome-wide topography of yeast meiotic recombination initiation. Cell 144: 719–731.

Peck JR. 1994. A ruby in the rubbish: Beneficial mutations, deleterious mutations and the evolution of sex. Genetics 137: 597–606.

Qi J, Chen Y, Copenhaver GP, and Ma H. 2014. Detection of genomic variations and dna polymorphisms and impact on analysis of meiotic recombination and genetic mapping. Proceedings of the National Academy of Sciences of the United States of America 111: 10007–10012.

Ritz KR, Noor MA, and Singh ND. 2017. Variation in recombination rate: Adaptive or not? Trends in Genetics 33: 364–374.

Rodgers-Melnick E, Bradbury PJ, Elshire RJ, Glaubitz JC, Acharya CB, Mitchell SE, Li C, Li Y, and Buckler ES. 2015. Recombination in diverse maize is stable, predictable, and associated with genetic load. Proceedings of the National Academy of Sciences of the United States of America 112: 3823–3828.

Rowan BA, Heavens D, Feuerborn TR, Tock AJ, Henderson IR, and Weigel D. 2019. An ultra high-density Arabidopsis thaliana crossover. Genetics 213: 771–787.

Saintenac C, Faure S, Remay A, Choulet F, Ravel C, Paux E, Balfourier F, Feuillet C, and Sourdille P. 2011. Variation in crossover rates across a 3-mb contig of bread wheat (triticum aestivum) reveals the presence of a meiotic recombination hotspot. Chromosoma 120: 185–198.

Salomé PA, Bomblies K, Fitz J, Laitinen RA, Warthmann N, Yant L, and Weigel D. 2012. The recombination landscape in Arabidopsis thaliana f2 populations. Heredity 108: 447–455.

Serra H, Choi K, Zhao X, Blackwell AR, Kim J, and Henderson IR. 2018. Interhomolog polymorphism shapes meiotic crossover within the Arabidopsis RAC1 and RPP13 disease resistance genes. PLoS Genetics 14.

Shen C, Wang N, Huang C, Wang M, Zhang X, and Lin Z. 2019. Population genomics reveals a fine-scale recombination landscape for genetic improvement of cotton. Plant Journal 99: 494–505.

Si W, Yuan Y, Huang J, Zhang X, Zhang Y, Zhang Y, Tian D, Wang C, Yang Y, and Yang S. 2015. Widely distributed hot and cold spots in meiotic recombination as shown by the sequencing of rice F2 plants. New Phytologist 206: 1491–1502.

Silva-Junior OB and Grattapaglia D. 2015. Genome-wide patterns of recombination, linkage disequilibrium and nucleotide diversity from pooled resequencing and single nucleotide polymorphism genotyping unlock the evolutionary history of Eucalyptus grandis. New Phytologist 208: 830–845.

Sun H, Rowan BA, Flood PJ, Brandt R, Fuss J, Hancock AM, Michelmore RW, Huettel B, and Schneeberger K. 2019. Linked-read sequencing of gametes allows efficient genome-wide analysis of meiotic recombination. Nature Communications 10: 1–9.

Sun Y, Ambrose JH, Haughey BS, Webster TD, Pierrie SN, Muñoz DF, Wellman EC, Cherian S, Lewis SM, Berchowitz LE, et al. 2012. Deep genome-wide measurement of meiotic gene conversion using tetrad analysis in Arabidopsis thaliana. PLoS Genetics 8: e1002968.

Szostak JW, Orr-Weaver TL, Rothstein RJ, and Stahl FW. 1983. The double-strand-break repair model for recombination. Cell 33: 25–35.

Wang J, Street NR, Scofield DG, and Ingvarsson PK. 2016. Natural selection and recombination rate variation shape nucleotide polymorphism across the genomes of three related populus species. Genetics 202: 1185–1200.

Weisweiler M, Arlt C, Wu PY, Inghelandt DV, Hartwig T, and Stich B. 2022. Structural variants in the barley gene pool: precision and sensitivity to detect them using short-read sequencing and their association with gene expression and phenotypic variation. Theoretical and Applied Genetics 135: 3511–3529.

Wijnker E, James GV, Ding J, Becker F, Klasen JR, Rawat V, Rowan BA, de Jong DF, de Snoo CB, Zapata L, et al. 2013. The genomic landscape of meiotic crossovers and gene conversions in Arabidopsis thaliana. eLife 2: e01426.

Yang S, Yuan Y, Wang L, Li J, Wang W, Liu H, Chen JQ, Hurst LD, and Tian D. 2012. Great majority of recombination events in Arabidopsis are gene conversion events. Proceedings of the National Academy of Sciences of the United States of America 109: 20992–20997.

Yelina NE, Choi K, Chelysheva L, Macaulay M, de Snoo B, Wijnker E, Miller N, Drouaud J, Grelon M, Copenhaver GP, et al. 2012. Epigenetic remodeling of meiotic crossover frequency in arabidopsis thaliana DNA methyltransferase mutants. PLoS Genetics 8: e1002844.

